# Synergism of gut microbiota to double-stranded RNAs in RNA interference of a leaf beetle

**DOI:** 10.1101/824581

**Authors:** Letian Xu, Shijing Xu, Liuwei Sun, Yiqiu Zhang, Jing Luo, Ralph Bock, Jiang Zhang

## Abstract

RNA interference (RNAi) has emerged as an efficient tool to control insect pests. When lethal double-stranded RNAs (dsRNAs) were ingested by the insects, strong gene silencing and mortality can be induced. To exert their function, dsRNA molecules must pass through insect’s gut and enter epithelial cells and/or the hemolymph. Gut bacteria are known to inhabit on the epithelial surface to confer host new capabilities to counter both biotic and abiotic stress. Whether there is a crosstalk between gut bacteria and dsRNAs and the effects of the microbiome on RNAi efficiency remains unknown. Here, using a leaf beetle-gut microbiota system, we investigated whether and how gut bacteria interact with dsRNA molecules and its effects on host insects. We firstly showed that the leaf beetle *Plagiodera versicolora* (Coleoptera) is highly susceptible to RNAi. Then, we found that ingestion of dsRNAs by non-axenic *P. versicolora* larvae results in (i) significantly accelerated mortality compared to axenic larvae, and (ii) over-growth and dysbiosis of the gut microbiota. The latter is mainly caused by the bacterial utilization of the dsRNA degraded products initiated by the host insect. Furthermore, we found that *Pseudomonas putida*, a gut bacterium of *P. versicolora*, was a main commensal-to-pathogen strain that accelerated the death of *P. versicolora* larvae. Taken together, our findings reveal a synergistic role of gut microbiota to dsRNA-induced mortality of pest insects, which provides new insights in the ecological functions of insect gut bacteria, and also contributes to a better understanding of the RNAi mechanisms in insects.

## Introduction

RNA interference (RNAi) is a conserved post-transcriptional gene-silencing process in which gene expression in eukaryotic cells is down-regulated by double-stranded RNAs (dsRNAs) and their degradation intermediates. RNAi has been widely used as a powerful experimental tool for determining gene functions, and more recently, was repurposed as a novel strategy for insect pest control. When dsRNAs that target essential insect genes are ingested by the pest insect, knock-down of the target genes by RNAi can result in reduced growth or even the death (1). RNAi-based pest control is, therefore, recognized as promising approach to develop new target-specific pesticides. Coleopteran insects, such as the red flour beetle (*Tribolium castaneum*), the western corn rootworm (*Diabrotica virgifera virgifera*), and the Colorado potato beetle (*Leptinotarsa decemlineata*, CPB) are highly susceptible to RNAi and can be efficiently killed, if lethal dsRNAs are orally delivered. Interestingly, the efficiency of RNAi-mediated pest control was shown to be highly variable among different groups of insects (2–5). A few factors influencing the RNAi efficiency in insects have been identified, including the stability of dsRNA in the digestive system, the cellular uptake efficiency of dsRNAs, endosomal entrapment, and the activity of the core RNAi machinery (6–8). However, a comprehensive understanding of the molecular mechanisms affecting RNAi efficiency is still lacking. Moreover, the complex network of molecular interactions underpinning the dsRNA-mediated killing mechanisms has received limited attention.

The gut of insects provides a distinctive and complex environment for microbial colonization. The gut microbiota of insects offers various beneficial services to the host, including multifarious contributions to food digestion, nutrient modification, pathogen defense, chemical detoxification, etc. (9). In addition to the positive contribution to the insect’s fitness, the presence of gut bacteria has also been implicated in promoting the insecticidal activity of *Bacillus thuringiensis* (10–13). Furthermore, it was demonstrated that some commensal gut bacteria have the potential to translocate from the gastrointestinal tract into the hemolymph and induce a rapid death (14–16). Upon ingestion of dsRNA by insects, the dsRNA molecules inevitably pass through the intestinal tract, where they get in contact with the diverse microbial community that thrives in the gut. Whether there is an interplay between the gut microbiota and exogenously applied dsRNAs, and whether or not the microbiome affects the RNAi response of host insect is largely unknown.

The willow leaf beetle, *Plagiodera versicolora*, is one of the most damaging pest species of Salicaceae plants such as willows and poplars (17). This pest is widely distributed across northern Africa, America, Europe and Asia. *P. versicolora* larvae and adults both feed on leaves of willows and poplars, especially during the summer months. If not controlled, the insects can completely skeletonize the leaves and cause serious economic damage (18).

In the present work, we have studied the response of *P. versicolora* to environmental RNAi. We found that, like many other Coleopteran insects, *P. versicolora* are susceptible to RNAi, and can be efficiently killed by feeding with dsRNAs against suitable essential genes. We further explored the role of the gut microbiota in the RNAi response in *P. versicolora*. Our results indicate that the gut microbiota can utilize the dsRNA degradation products for proliferation, and contribute substantially to insect mortality.

## Materials and methods

### Preparation of *P. versicolora* larvae and gut bacterial strains

*P. versicolora* insects used in the studies were reared by feeding with detached fresh willow leaves or axenic poplar leaves at 28 °C and 60% ± 5% relative humidity under a 16 h light /8 h dark photoperiod. Poplar plantlets was raised aseptically on a synthetic medium containing 4.4 g/L MS salts plus vitamins, 0.1 mg/L NAA and 3 % (w/v) sucrose (19). To obtain axenic larvae, egg masses were soaked in 40% NaOH for 1 min and subsequently in 70% ethanol for 5 min, and finally washed with sterilized water for 2 min. After air-drying, each egg was separately transferred onto Luria-Bertani (LB) solid medium. Removal of gut bacteria was confirmed by a colony forming unit assay and PCR analysis using conserved primers for the 16S rRNA gene of gut bacteria. The newly hatched larvae were fed with aseptic detached poplar leaves and maintained axenically.

Strains of the gut bacteria *Enterobacter aerogenes, Pseudomonas putida* and *Enterococcus faecalis* strains were isolated from *P. versicolora*. To identify the bacterial species, samples of genomic DNA from the different bacterial isolates were obtained by extraction with the High Pure PCR Template preparation kit (Roche), and used to amplify 16S rRNA gene sequences by PCR using the universal 16S rRNA gene-specific primers 8F and 1492R (Table S1). The purified PCR products were then sequenced (Sangon Biotech, Shanghai), and the obtained sequences were aligned with the closest relatives matching the 16S rRNA gene sequences by BLAST searches (http://blast.ncbi.nlm.nih.gov/Blast.cgi).

### Ingestion and injection of dsRNA in *P. versicolora* larvae

*Srp54k, Actin, Snap, Shi*, and *EGFP* gene fragments were amplified by PCR using gene-specific primer pairs (Table S1), and subsequently used as templates to synthesize dsRNA (ds*Srp54k*, ds*Actin*, ds*Snap*, ds*Shi*, and ds*GFP*) with the T7 RiboMAX^TM^ Express RNAi system (Promega, USA) following the manufacturer’s instructions. Integrity of dsRNAs was evaluated by electrophoresis in 1.5% agarose gels and the amounts of dsRNA were quantified with a spectrophotometer (Nano-Drop 2000, Thermo Scientific, USA).

For dsRNA feeding assays, 30 second instar larvae were fed with aseptic poplar leaves that had been painted with 8 ng/cm^2^ dsRNA. Survival was recorded daily. To examine the efficiency of RNAi-mediated knock-down of target genes, five larvae were randomly chosen each day after dsRNA feeding and total RNA was isolated for further analysis.

For dsRNA injection assays, 60 second instar larvae were anaesthetized on ice before injection, and samples of 36 ng ds*Srp54k* or 2 ng ds*Actin* in 5 nL were injected into each larva using a micro-injector (World Precision Instruments, Sarasota, USA). Negative controls received an equivalent amount of ds*GFP*. After injection, *P. versicolora* larvae were reared on sterile poplar leaves, and survival rates were monitored daily (n = 30). Five larvae were randomly chosen each day for RNA extraction.

To suppress dsRNase activity, larvae were fed with fresh poplar leaves painted with 40 ng/cm^2^ ds*dsRNase* and 8 ng/cm^2^ ds*GFP* or, as a control, the equivalent amount of ds*GFP* (48 ng/cm^2^). Five larvae were randomly chosen after four days of feeding for RNA extraction.

### Analysis of gut microbiota

After three days of dsRNA feeding (including H_2_O as a negative control), larvae were disinfected in 70% ethanol for 10 s, rinsed in sterile water several times, and then dissected under sterile conditions. Four guts were pooled together as a replicate (n = 5). DNA was extracted from gut samples with the High pure PCR template preparation kit (Roche). A 16S rRNA gene fragment encompassing the V3 and V4 hypervariable regions was amplified by PCR using the primer pair 338F and 806R (Table S1).

Sequencing of 16S rRNA amplicons was performed using an Illumina MiSeq platform in MAJORBIO. Sequences were assigned to samples according to specific barcodes, followed by removal of barcode and primer sequences. Paired-end reads were assemble using FLASH (V1.2.7). High-quality data (clean reads) were obtained by using the QIIME (Quantitative Insights Into Microbial Ecology) software packages (V1.9.0) by filtering low-quality data with default parameters. Chimeric sequences were checked and removed using the UCHIME algorithm. All effective reads from each sample were initially clustered into Operational Taxonomic Units (OTUs) of 97% sequence similarity with a UPARSE algorithm. The most abundant sequence in each OTU was selected as the representative OTU and annotated by the RDP classifier algorithm and reference data sets from the SILVA database under a confidence threshold of 70%.

### *In vitro* incubation of dsRNA with hemolymph or gut juice

Hemolymph fluid samples were collected with a 10 μL glass capillary tube from abdomen of 10 fourth-instar larvae and were subsequently diluted with 100 μL ice-cold Ringer solution (1 L: 8.766 g NaCl; 0.188 g CaCl2; 0.746 g KCl; 0.407 g MgCl2; 0.336 g NaHCO3; 30.807 g sucrose; 1.892 g trehalose; pH 7.2) (20). Samples were then centrifuged at 12,000 *g* for 10 min to remove hemocytes and the supernatant was collected. For gut juice preparation, whole guts from 10 fourth-instar larvae were dissected and then crushed in 100 μL ice-cold Ringer solution in a centrifuge tube, followed by centrifugation at 12,000 *g* for 10 min and collection of the supernatants. The total protein concentrations of the fluid samples were determined using the BCA Protein Assay kit following the manufacturer’s instructions. 150 ng ds*GFP* (dissolved in 5 μL nuclease-free water) were added to 5 μL hemolymph or gut juice and incubated at 30 °*C* for the indicated times. Incubation in Ringer solution with the same amount of ds*GFP* served as control. Subsequently, 2 μL of 6× loading dye (20% glycerol, 1.25mM Na2ETDA, 0.1% bromophenol blue, 0.1% xylene cyanol) was applied to the 10 μL samples, and the RNAs were analyzed by electrophoreses in 1% agarose gels containing Gelview.

### Bacterial growth promotion and inhibition assays

To measure the effects of dsRNA degradation products on growth of gut bacteria, bacterial cultures were grown in a modified minimal medium (21). For carbon source determination, bacteria were grown in minimal medium supplemented with 24 nmol/mL ds*GFP*, NTP mix, uridine or inosine as sole carbon source and 0.4% (NH4)2SO4 as nitrogen source. For nitrogen source determination, bacteria were grown in minimal medium supplemented with 4% glucose and 0.05% sodium citrate as carbon source and 24 nmol/mL ds*GFP*, NTP mix, uridine or inosine as sole nitrogen source. Saline-washed cells (approximately 10^7^ cells) of each bacterial strain were used for inoculation. Subsequently, the cultures were shaken for 24 h at 200 rpm and 30 °C. The bacterial cell number was determined by counting colony-forming units (CFUs) on LB medium.

### Reintroduction of bacteria into beetle guts

After overnight culture, bacteria were collected by centrifugation at 4,000 rpm for 5 min to remove the supernatant. The bacterial cells were then washed with sterile PBS 3 times, re-suspended in PBS, and diluted to a final concentration of approximately 1× 10^6^ cells/mL (22). The bacterial suspension was mixed with the indicated amounts of dsRNA and painted onto aseptic poplar leaves. The leaves were then fed to axenic *P. versicolora* larvae (n = 40). The aseptic poplar leaves coated with bacteria (approximately 1 × 10^4^ cells/cm^2^) and dsRNA (8 ng/cm^2^) were replaced every day. Survival was recorded daily.

### Quantitative real-time PCR analysis

RNA was extracted using RNAiso plus as described by the manufacturer (Takara, Japan). cDNA was synthesized from 2 μg of total RNA using the Hifair^@^ 1^st^ Strand cDNA Synthesis for qPCR Kit (YEASEN) with random primers. qPCRs were performed in an Applied Biosystems 7300 Real-Time PCR system using SYBR premix Ex Taq^TM^ (Takara, Japan). cDNA templates were denatured at 95°C for 2 min, followed by 40 two-segment cycles of amplification at 95°C (5 s) and 60°C (30 s), where the fluorescence was automatically measured. A melting curve analysis was performed after the qPCR run (between 65°C and 95°C with 0.5°C increments). Prior to use in qRT-PCR, cDNA was 1:9 diluted with H_2_O. All data were normalized to levels of a housekeeping gene (5S rRNA gene), and gene expression levels were calculated as fold change values using the 2^-ΔΔCt^ method (23). Each experiment was carried out in triplicate. All primers used were designed by Primer Premier 6. Primer sequences are listed in Table S1.

### Histological analysis

Larval tissues were immediately fixed in 10% neutral buffered formalin supplemented with 2% dimethyl sulfoxide for 24 h, then dehydrated in a series of alcohol baths (beginning with 50% and progressing to 100%), cleared in xylol for 4 h, and finally embedded in paraffin. Cross-sections were prepared with a microtome (LEICA RM 2016, Leica Microsystems, USA) at a thickness of about 3 μm, stained with hematoxylin and eosin, and analyzed with a fluorescence microscope (Nikon Eclipse E-200 model, Tokyo, Japan).

### Statistical analysis of data

Data comprising two groups were analyzed using Student’s *t* test for unpaired comparisons, and data comprising more than two groups were analyzed using one-way ANOVA coupled with Bonferroni (equal variances) or Dunnett’s T3 (unequal variances) correction multiple comparison test. Survival curves were analyzed using the Kaplan– Meier method, and the log-rank test was used to evaluate the significance of differences between two groups. A value of P<0.05 was considered significantly different. Data were statistically analyzed using SPSS version 19.0. Figures were drawn using Origin 8.5 and GraphPad Prism 7, and assembled in Adobe Illustrator CS6 and Photoshop CS6.

## Results

### Gut microbiota accelerate dsRNA-induced mortality of *P. versicolora* larvae

We first wanted to examine the sensitivity of *P. versicolora* to RNAi and identify essential genes that represent suitable targets for RNAi-mediated pest control (24). To this end, larvae were fed with *in vitro* synthetsized dsRNAs targeted against the *β-Actin*, *Srp54k*, *Snap* and *Shi* genes. *β-Actin* encodes a multi-functional protein for microfilament formation (25). *Srp54k* encodes the 54 kDa subunit of the signal recognition particle which is involved in protein targeting to the endoplasmic reticulum (26). *Snap* encodes the alpha-soluble N-ethylmaleimide-sensitive factor attachment protein that is involved in the docking and fusion of vesicles to target membranes (27). *Shi* encodes a dynamin-like protein associated with vesicular trafficking (28). Compared to the negative control (feeding with *GFP* gene-derived dsRNA; ds*GFP*), feeding of *P. versicolora* with dsRNAs derived from *Srp54k, Actin* and *Snap* resulted in 100 % mortality of the larvae. The effects of RNAi were shown to be ds*Actin* > ds*Srp54k* >ds*Snap* (Fig. S1; log-rank test, *P* < 0.05).

To evaluate whether the gut microbiota of *P. versicolora* is involved in determining the efficiency of dsRNA-mediated killing, axenic larvae were obtained by surface sterilization of eggs. The hatched larvae were then reared on aseptically grown poplar leaves. The axenic status of the larvae was confirmed by two independent methods: (i) absence of bacterial colony formation on LB agar plates, and (ii) absence of PCR products from amplification reactions using primers for conserved regions of the bacterial 16S rDNA (Fig. S2a,b). The axenic growth had no influence on insect survival compared to conventionally raised larvae (Fig. S2c).

dsRNA feeding assays revealed no significant differences in survival of axenic larvae compared to non-axenic larvae when fed with ds*GFP*. By contrast, non-axenic larvae fed with ds*Srp54k* or ds*Actin* were killed significantly faster than axenic larvae (Fig. 1a,c; log-rank test, *P*<0.001). Surprisingly, no significant differences in the level of gene silencing were found between axenic and non-axenic larvae (Fig. 1b,d; log-rank test, *P*<0.05). This finding suggests that the striking differences in mortality between non-axenic and axenic larvae fed with lethal dsRNAs are not caused by differences in the strength of the suppression of the targeted genes. Instead, these results raise the interesting possibility that the gut microbiota confers the accelerated dsRNA-mediated mortality in non-axenic *P. versicolora* larvae.

**Fig. 1.**
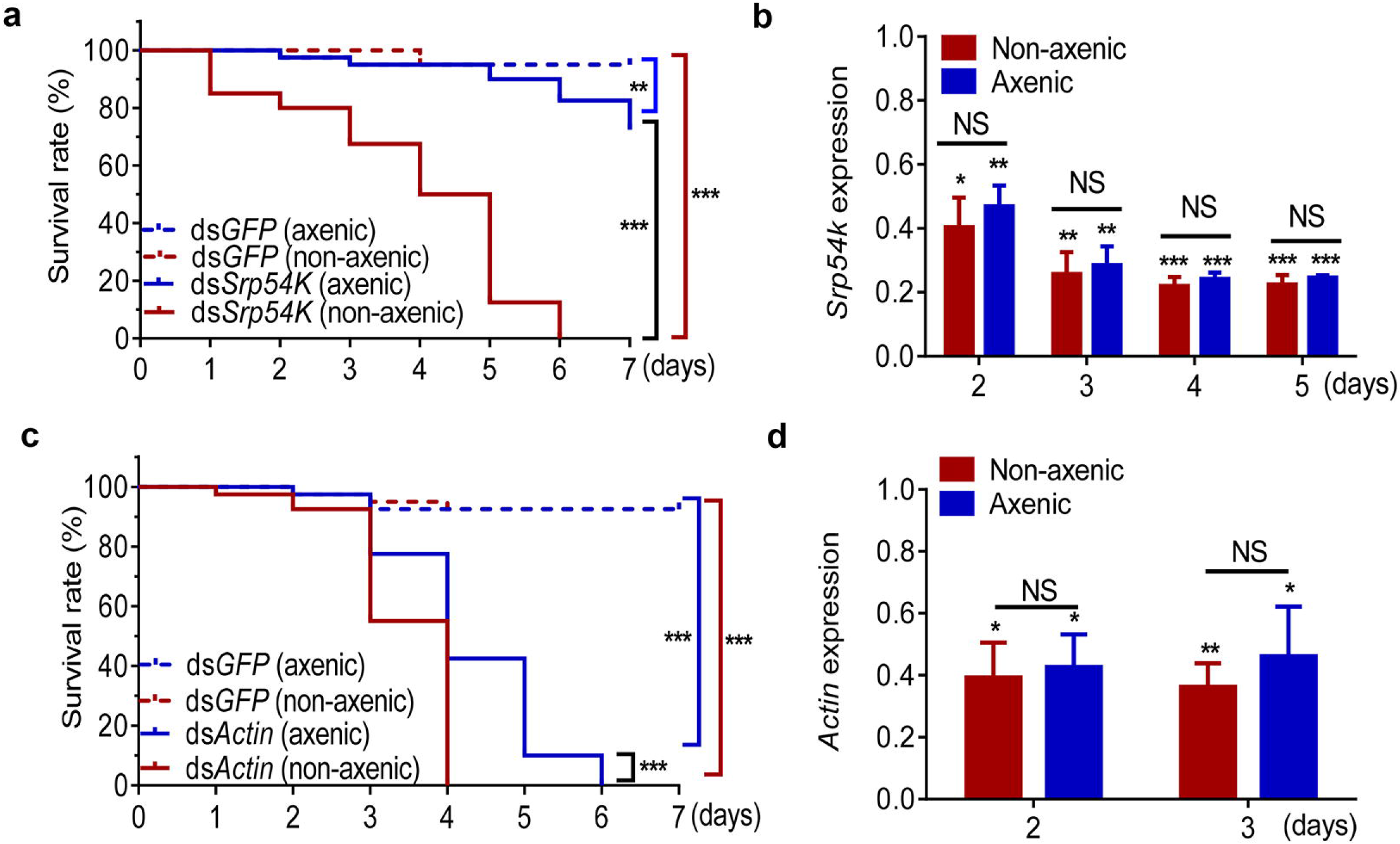
Gut bacteria accelerate mortality of *P. versicolora* in lethal dsRNA-mediated killing process. (**a** and **c**) Kaplan-Meier survival curves of axenic and non-axenic larvae (n=30) after feeding with ds*Srp54k* (**a**) and ds*Actin* (**c**). *Srp54k* (**b**) and *Actin* (**d**) relative expression in axenic and non-axenic larvae upon fed on ds*Srp54k* and ds*Actin*. Gene expression levels were set as one in larvae fed with ds*GFP* control. Kaplan-Meier survival curve was analyzed by the log-rank test. Gene expression differences between two groups were calculated using the independent-samples *t*-test. Data are presented as mean ± SE, *** *P* < 0.001; ** *P* < 0.01; * *P* < 0.05; NS, not significant.

To examine whether breakdown of the barrier function of the gut epithelium is responsible for the enhanced killing efficiency, we isolated hemolymph from non-axenic larvae fed with dsRNAs and assayed the hemolymph samples for the presence of bacteria by culturing them on LB medium. While numerous bacterial colonies formed from non-axenic larvae fed with ds*Srp54k* or ds*Actin*, no colonies grew from hemolymph samples obtained from larvae fed with ds*GFP*. This finding indicates translocation of gut bacteria to the hemocoel in *P. versicolora* larvae fed with lethal dsRNAs (ds*Srp54k* or ds*Actin*; Fig. S3). Moreover, histological analysis revealed that the gut epithelia of non-axenic larvae fed with lethal dsRNAs were disrupted, while they remained intact in the ds*GFP* control (Fig. S4). These observations provide further evidence for gut bacteria promoting dsRNA-mediated killing.

### Feeding of dsRNAs results in alteration of the composition of the gut microbiota of *P. versicolora*

We next investigated the composition of the gut bacterial community in *P. versicolora* and its change after ingestion of lethal dsRNAs (ds*Srp54k* or ds*Actin*) or ds*GFP* as control. Ingestion of sterilized water served as negative control. 878,019 sequences (96.5% of the total number of 909,726 trimmed reads) were obtained from 16S rDNA amplicon sequencing, and grouped into 122 operational taxonomic units (OTUs) using a 97% similarity cut-off level. Alpha diversity was estimated using four measurements: ACE index, Chao1 index, Shannon–Weiner index, and Simpson’s index (Table S2). In general, no significant differences were found for the α-diversity indexes between gut samples of control animals and the three groups of larvae feeding on dsRNA-painted leaves. However, the larvae fed with ds*Actin* had a significantly greater value for the Shannon diversity index and a significantly lower Simpson’s index than the other treatments (Table S2; one-way ANOVA, *P*<0.05), possibly suggesting that the oral administration of ds*Actin* can increase bacterial community evenness.

Principal co-ordinates analysis (PCoA) of Bray-Curtis distances of microbial communities showed that bacterial communities from the control larvae clustered independently and distinctly from the three dsRNA-fed larvae (Fig. 2a; PERMANOVA, *P* < 0.05). Typing analysis of the four treatments indicated that they fall into three major types (Fig. 2b). In all samples, the gut bacterial community was dominated by four bacterial genera: *Enterobacter*, *Pseudomonas*, *Enterococcus* and *Serratia*, which account for over 95% of the total sequences in the samples (Fig. 2c; Table S3). The proportion of *Enterobacter* was significantly reduced in the three groups of dsRNA-fed larvae compared to the control group (larvae feeding on dsRNA-free leaves) (Fig. 2c; Kruskal-Wallis *H* test, *P* < 0.05), and the proportion of two major genera (*Pseudomonas* and *Enterococcus*) increased after feeding with the dsRNAs. Remarkably, feeding with lethal dsRNAs (i.e., ds*Srp54k* or ds*Actin*) significantly increased the proportion of *Pseudomonas* in *P. versicolora* larvae compared to larvae fed with untreated or ds*GFP*-treated leaves (Fig. 2c; Kruskal-Wallis *H* test, *P* < 0.05).

**Fig. 2.**
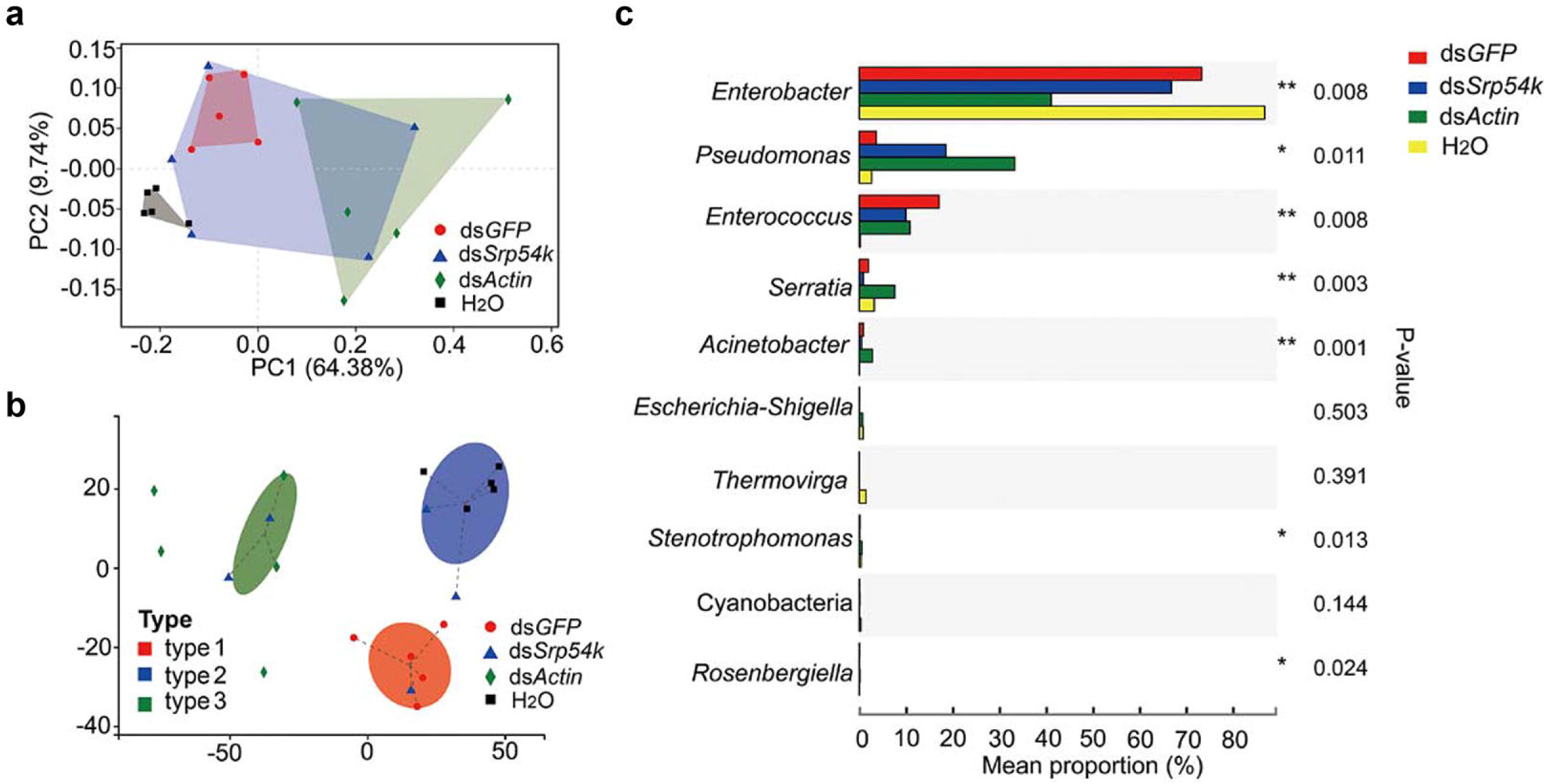
Ingestion of dsRNA alters gut microbiota composition of *P. versicolora*. (**a**) Principal co-ordinates analysis (PCoA) of Bray-Curtis distances at the genus level for 16S rRNA gene sequences. (**b**) Typing analysis of Jensen-Shannon divergence distance at the OTU level for 16S rRNA gene sequence data. (**c**) Kruskal-Wallis *H* test bar plot analysis of the relative proportions of major genera of gut bacteria. The relative abundances of the identified microbial taxa were determined in gut samples collected from larvae fed with ds*GFP,* ds*Srp54k*, ds*Actin*, or H_2_O. *P*-values were calculated using the Kruskal-Wallis *H* test. ** *P* < 0.01; * *P* < 0.05.

### Ingestion of dsRNA promotes growth of gut bacteria

The total amounts of gut bacteria were significantly increased in all dsRNA-fed larvae (2 days after feeding with ds*GFP* and ds*Actin*, 3 days after feeding with ds*Srp54k*) compared to the control group of larvae feeding on leaves not painted with dsRNA (Fig. 3a-c; *T*-test, *P* < 0.05). This finding suggests that ingestion of dsRNA can promote bacterial growth in the intestinal system. To further validate the effects of dsRNA ingestion on the abundance of the three major genera of gut bacteria, we monitored the relative quantities of the bacterial genera in the different groups of larvae. The abundances of *Enterobacter* and *Pseudomonas* were found to be significantly increased in larvae fed with ds*GFP*-treated leaves at day 3 and 4 compared to the control (Fig. 3d; *T*-test, *P* < 0.05), but dropped to similar abundance at day 5. The relative abundances of both *Pseudomonas* and *Enterobacter* were significantly increased after 3 days of feeding on ds*Srp54k*, while no significant differences were found in relative abundance of *Enterococcus* between the larvae fed with ds*Srp54k* and the control (Fig. 3e; *T*-test, *P* < 0.05). Similarly, the relative abundances of both *Pseudomonas* and *Enterococcus* were significantly increased after 3 days of ds*Actin* feeding (Fig. 3f; *T*-test, *P* < 0.05).

**Fig. 3.**
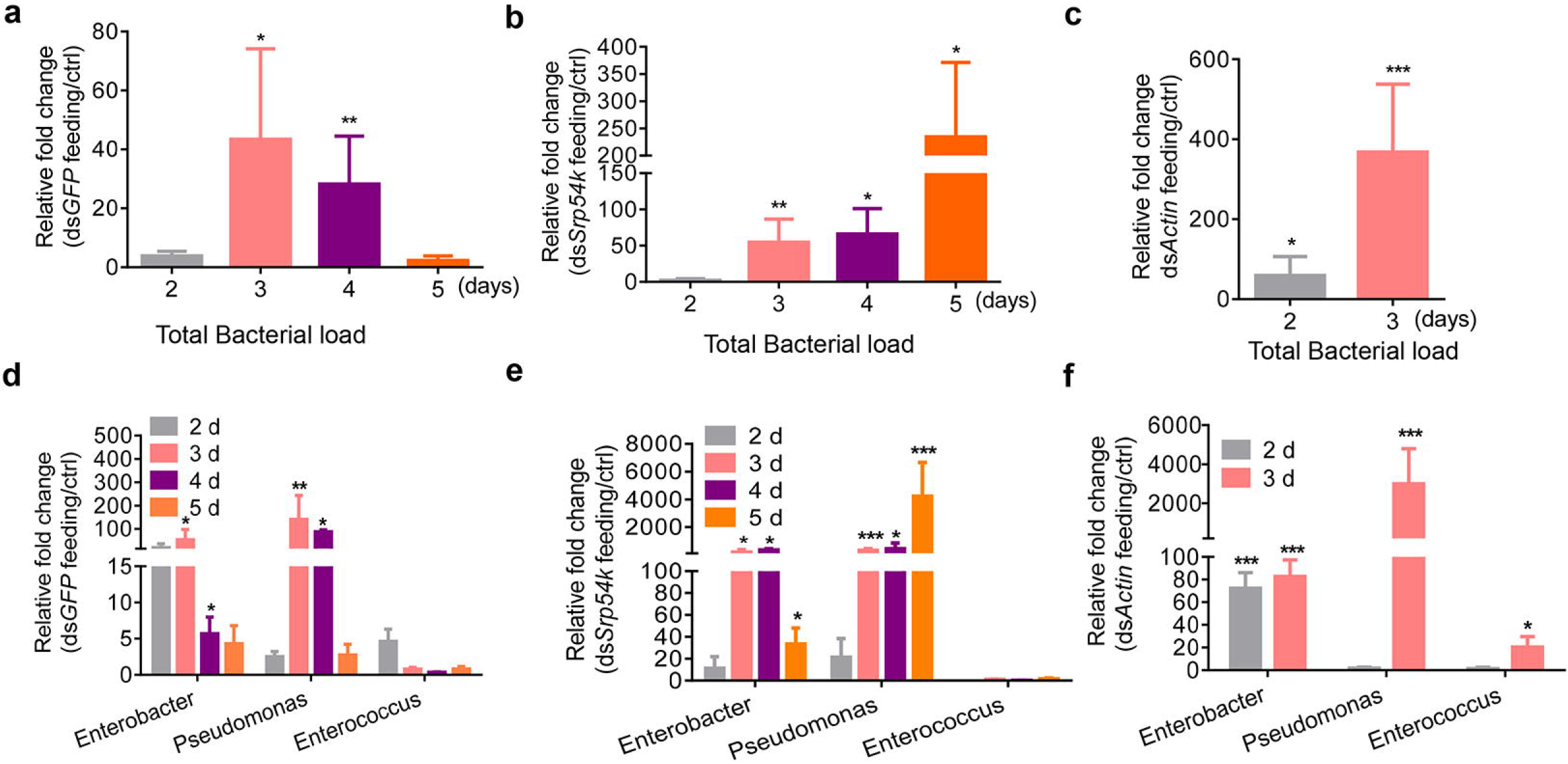
Ingestion of dsRNA promotes the growth of gut bacteria in *P. versicolora* larvae. qRT-PCR analyses were performed to determine the relative abundance of gut bacteria in *P. versicolora* larvae fed with ds*GFP* (**a**), ds*Srp54k* (**b**) and ds*Actin* (**c**) (n = 5), and the abundances of three major bacterial genera in non-axenic *P. versicolora* larvae fed with ds*GFP* (**d**), ds*Srp54k* (**e**) and ds*Actin* (**f**). The measurements were performed at different time points (after 2-5 days of feeding). The qRT-PCR value obtained for gut bacterial 16S rRNA in *P. versicolora* control larvae (fed with water-treated leaves) was set as 1.0. *P*-values were calculated using the independent-samples *t*-test. Data are presented as means ± SE, *** *P* < 0.001; ** *P* < 0.01; * *P* < 0.05; NS, not significant.

### Ingestion of dsRNA is confers accelerated mortality and increased gut bacterial load in *P. versicolora*

To assess the contribution of ingestion of insecticidal dsRNA to the proliferation of gut bacteria, we determined mortality and bacterial loads of *P. versicolora* larvae after feeding and injection of dsRNAs. Upon injection of dsRNA, a significantly accelerated mortality was observed in larvae treated with ds*Srp54k* compared to those injected with ds*GFP* (Fig. 4a; log-rank test, *P* < 0.05). Moreover, we found that non-axenic larvae fed with ds*Srp54k* exhibit higher mortality and gut bacterial loads than larvae injected with ds*Srp54k* (Fig. 4a; log-rank test, *P* < 0.05; Fig. 4c, *T*-test, *P* < 0.05), although the silencing levels of the target gene (*Srp54k*) were similar in the two treatments (Fig. 4b). Ingestion of ds*Actin* by *P. versicolora* larvae lead to a similar morality as injection of ds*Actin* (Fig. 4d), even though the latter treatment caused a significantly higher efficiency of gene silencing (Fig. 4e; *T*-test, *P* < 0.05) and a lower gut bacterial load (Fig. 4f, *T*-test, *P* < 0.05). Taken together, these results indicate that ingestion of insecticidal dsRNA accelerates mortality of *P. versicolora* larvae.

**Fig. 4.**
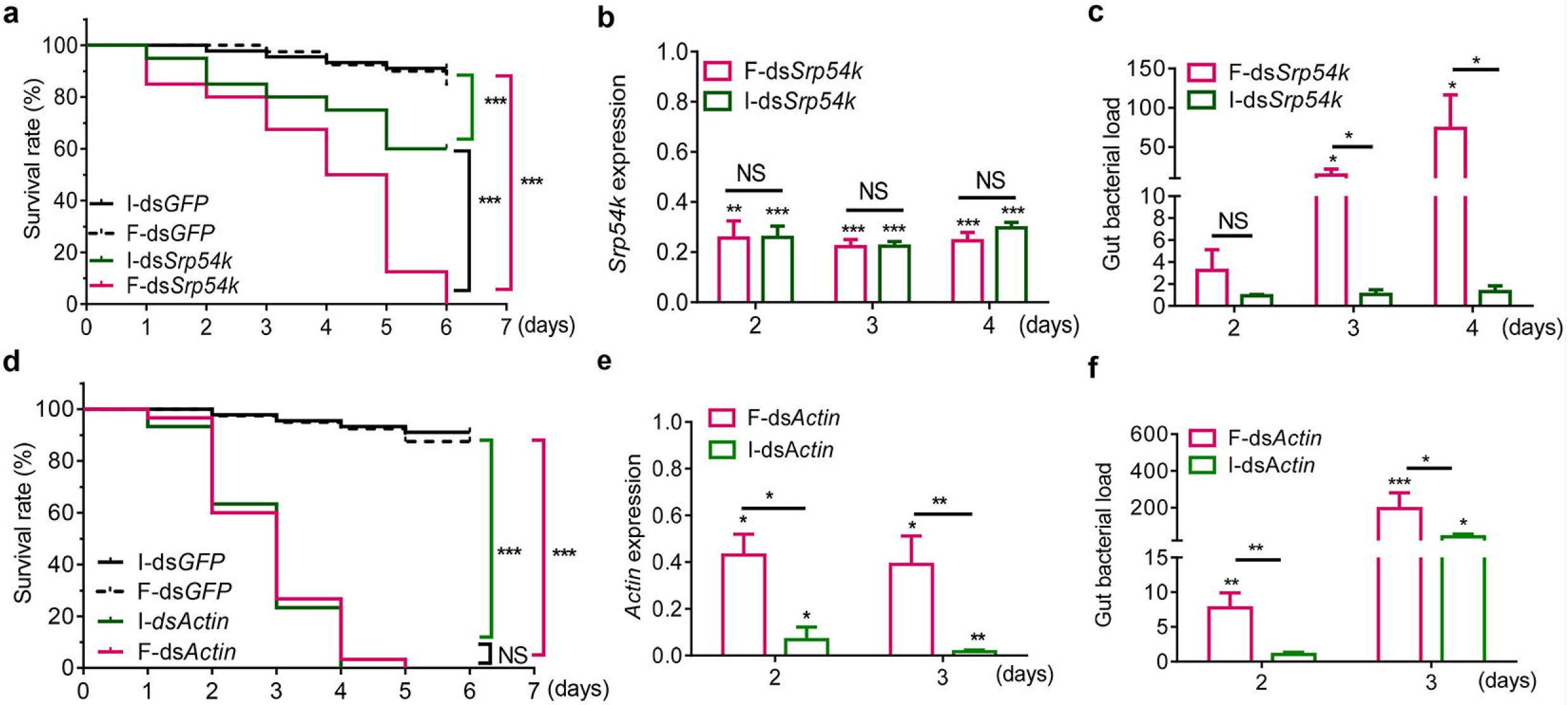
Comparison of mortality of *P. versicolora* in response to insecticidal dsRNA upon administration by injection (injected dsRNA, I-dsRNA) or ingestion (fed dsRNA, F-dsRNA). (**a, d**) Kaplan–Meier survival curves of *P. versicolora* larvae (n = 30) after injection and ingestion of ds*Srp54k* (**a**) or ds*Actin* (**d**). Ingestion or injection of ds*GFP* were performed as controls. The survival curves were analyzed by the log-rank test. (**b, e**) Expression levels of *Srp54k* and *Actin* in *P. versicolora* larvae fed or injected with ds*Srp54k* (**b**) or ds*Actin* (**e**). Relative gene expression levels were determined by qRT-PCR assays and expression in control larvae treated with ds*GFP* were set as one. (**c,f**) qRT-PCR analysis of the relative abundance of whole gut bacteria in *P. versicolora* larvae after injection with ds*Srp54k* (**c**) or ds*Actin* (**f**) compared to ds*GFP* as control. *P*-values were calculated using the independent-samples *t*-test. *** *P* < 0.001; ** *P* < 0.01; * *P* < 0.05; NS, not significant.

### dsRNA instability in *P. versicolora* gut juice

To obtain information on the stability of dsRNA in the *P. versicolora* digestive system and the hemolymph, ds*GFP* was incubated in gut juice or hemolymph extracted from insects. While ds*GFP* was rapidly degraded in the gut juice, it remained relatively stable in the hemolymph (Fig. 5a). To determine the possible cause of dsRNA instability in the digestive tract, we retrieved two putative dsRNase sequences from our *P. versicolora* RNA-seq datasets. Both sequences have high sequence similarities to previously identified gut dsRNases (Fig. S5). *dsRNase1* was found to be highly expressed in the gut tissue compared to the carcass, while expression of *dsRNase2* was relatively low in both gut and carcass (Fig. 5b). Expression of *dsRNase1* is induced upon feeding of *P. versicolora* larvae with ds*GFP* (Fig. 5c). Importantly, the degradation capability of dsRNA by the gut juice of *P. versicolora* was strongly reduced when *dsRNase1* was down-regulated by injection of double-stranded RNA (ds*dsRNase1*) (Fig. 5d,e). These results indicate that dsRNase1 is the major dsRNA-degrading enzyme activity in the gut juice of *P. versicolora*.

**Fig. 5.**
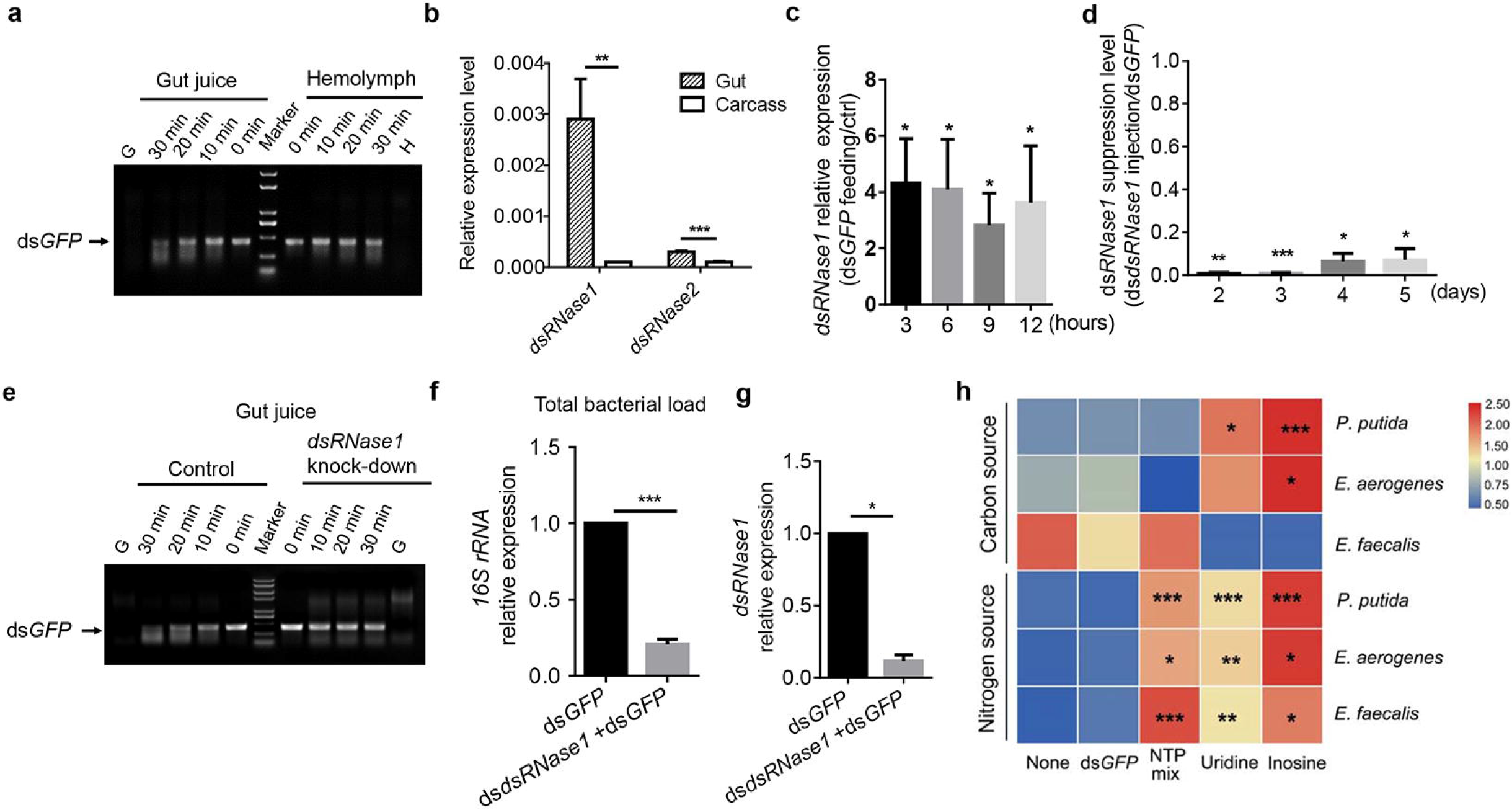
Degradation of dsRNA promotes gut bacterial growth in *P. versicolora* larvae. (**a**) ds*GFP* incubation with hemolymph or gut fluid of *P. versicolora* third instar larvae. Samples of 5 μL gut juice or hemolymph (adjusted to 5 μg of total protein) were incubated with 400 ng of *dsGFP* for the indicated times at 28 °C. G: gut juice sample without dsRNA added; H: hemolymph sample without dsRNA added. (**b**) Expression patterns of two putative *dsRNase* genes in third instar larvae. Each sample comprises five pooled guts or carcasses from dissected larvae. Error bars represent the mean of five independent samples ± SEM. (**c**) Relative expression changes of *dsRNase1* in larvae fed with ds*GFP*-painted leaves (8 ng/cm^2^) at the indicated times compared to untreated control larvae. Gene expression levels were set to 1 in larvae fed with H_2_O-treated control leaves (ctrl). (**d**) ds*RNase1* silencing efficiency over time in larvae (n = 5) injected with 40 ng of ds*dsRNase*. Injection of larvae with the same amount of ds*GFP* served as control. (**e**) ds*GFP* incubation in gut fluid of *P. versicolora* larvae with or without suppression of *dsRNase1* expression. Samples of 5 μL gut fluid (adjusted to 5 μg of total protein) were incubated with 400 ng of ds*GFP* for the indicated times at 28 °C. (**f**) qRT-PCR analysis of the relative abundance of gut bacteria in *P. versicolora* second instar larvae fed on young poplar leaves painted with ds*dsRNase1* (40 ng/cm^2^) and ds*GFP* (8 ng/cm^2^) or ds*GFP* (48 ng/cm^2^). Guts were sampled at day 4 (n = 7). (**g**) Relative expression levels of *d*s*RNase1* in the larvae assayed in (f). (**h**) dsRNA degradation products promote the growth of three gut bacterial species. Bacteria were grown for 24 h in either carbon-free or nitrogen-free liquid medium supplemented with 24 nmol/mL of the indicated compounds (dsGFP, NTP mix, inosine, uridine). Glucose (0.4%) and sodium citrate (0.05%) served as carbon source in nitrogen-free media. (NH_4_)_2_SO_4_ (0.4%) served as nitrogen source in carbon-free media. *** *P* < 0.001, ** *P* < 0.01, * *P* < 0.05.

### The degradation products of dsRNA can be utilized by gut bacteria for growth

To explore whether the instability of dsRNA contributes to the increased gut bacterial load, we fed larvae with dsRNA targeted against *dsRNase1* (ds*dsRNase1*) and dsGFP. For comparison, dsGFP was fed as a control. Interestingly, the gut bacterial load was significanly lower in the *dsRNase1*-silenced larvae than in larvae fed only with ds*GFP* (Fig. 5f,g), suggesting that dsRNA degradation is associated with increased gut bacterial load. To futher assess whether dsRNA or its degradation products promote gut bacterial growth, we isolated and identified three bacterial species (*Enterococcus aerogenes, P. putida,* and *E. faecalis*) from the gut of *P. versicolora*, and used ds*GFP* and RNA degradation products or building blocks (NTP mix, uridine, inosine) as either the sole carbon source or the sole nitrogen source for the growth of these bacterial species. As expected, ds*GFP* does not promote growth of any of the tested bacterial species. By contrast, all three species can utilize NTP mix, uridine and inosine as sole nitrogen source. The growth of *P. putida* and *E. aerogenes* (but not *E. faecalis*) also can be stimulated with uridine or inosine as sole carbon source (Fig. 5h; Table S3).

### Reintroduction of gut bacteria into axenic *P. versicolora* larvae enhances mortality upon administration of insecticidal dsRNA

To ultimately confirm the impact of gut bacteria on dsRNA-induced killing, individual gut bacterial species (*P. putida, E. aerogenes* or *E. faecalis*) were mixed with insecticidal dsRNA and reintroduced into axenic larvae. Compared to axenic larvae, mortality was significantly increased in larvae inoculated with bacteria. Upon feeding of ds*Srp54k*, *P. putida* inoculated larvae were killed significantly faster than larvae inoculated with *E. aerogenes* or *E. faecalis* (Fig. 6a; log-rank test, *P* < 0.05). Similar results were obtained for ds*Actin* feeding, and the accelerating effect of the gut bacteria was *P. putida* > *E. faecalis* > *E. aerogenes* (Fig. 6b; log-rank test, *P* < 0.05).

**Fig. 6.**
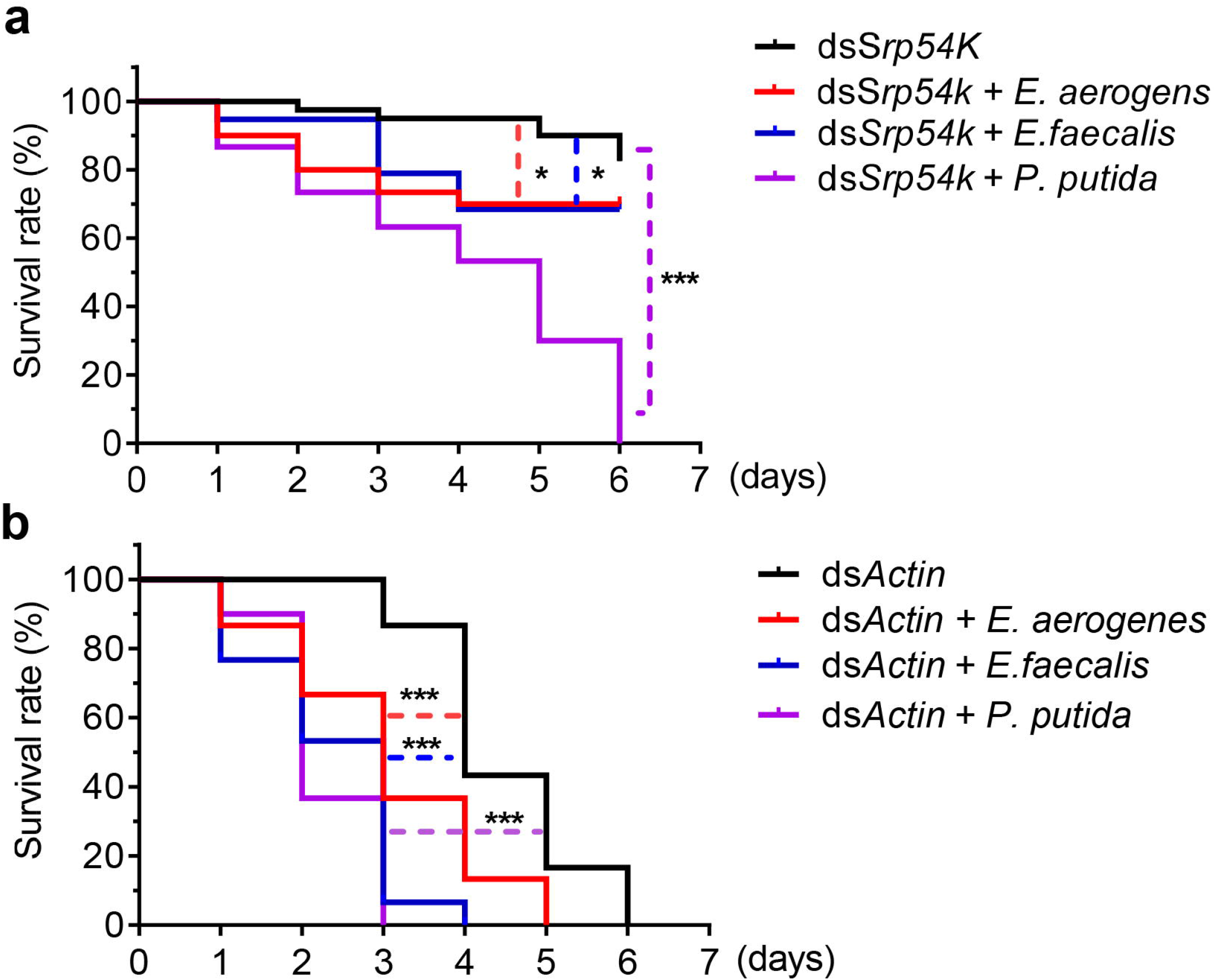
Kaplan–Meier survival curves of axenic *P. versicolora* larvae (n = 30) fed with ds*Srp54k* (**a**) or ds*Actin* (**b**) after reintroduction of bacterial species (*E. aerogenes*, *E. faecalis* or *P. putida*). The bacteria were mixed with either ds*Srp54k* or ds*Actin* and introduced into axenic larvae via spreading the bacteria onto aseptic poplar leaves. The log-rank test was used to assess the significance of differences between two survival curves. \*\*\**P* < 0.001, * *P* < 0.05.

## Discussion

In this work, we explored whether the gut microbiota plays a role in dsRNA-mediated killing of pest insects by RNA interference (environmental RNAi). To this end, we developed a method to obtain axenic larvae. The method uses surface sterilization of eggs, which has several advantages compared to the antibiotic treatments that are often used in microbiome research to eliminate bacteria, including complete removal of microbes and absence of residual antibiotics that may affect larval development upon hatching. The axenic status can be maintained by feeding with aseptic leaves of poplar plants grown in a sterile envirionment. Feeding dsRNAs targeted against two essential genes (*β-Actin* and *Srp54k*) to axenic and non-axenic larvae of *P. versicolora*, we found that ds*Actin* is more effective in killing *P. versicolora* than ds*Srp54k* in either axenic or non-axenic condition. The different killing efficiency might due the different cellular functions of the two proteins and/or differences in expression levels or transcript susceptibility to RNAi. Importantly, regardless of which dsRNA was fed, non-axenic larvae were killed significantly faster than axenic larvae. This effect was independent of the efficiency of gene knock-down (Fig. 1), excluding the possibility that the different mortalities between axenic and non-axenic larvae are caused by different levels of gene silencing of the targeted genes.

To pinpoint potential causes of the accelerated mortality of non-axenic larvae, we determined abundance and composition of the gut microbiota in larvae after ingestion of dsRNA. Compared to control larvae, feeding of dsRNAs (irrespective of RNAi induction and target sequences, i.e., feeding with ds*GFP*, ds*Actin* or ds*Srp54k*) resulted in significant alteration of the composition of the gut microbiota (Fig. 2) and excessive growth of gut bacteria (Fig. 3).

The most abundant bacterial genus in the gut microbiota is *Enterobacter*, comprising over 80% of the total microbiota (Fig. 2c). In previous studies, *Enterobacter* was also found to be widely associated with leaf beetles and poplar pests (29, 30). Bacteria of the genus *Rahnella* also could be readily isolated from *P. versicolora* using culture-dependent methods (18). *Enterobacter* from the Colorado potato beetle was shown to suppress plant defense (30); however, whether *Enterobacter* in *P. versicolora* has a similar function remains to be investigated. Notably, we found that there was a drastic increase in the abundance of *Pseudomonas* in larvae fed with ds*Srp54k* or ds*Actin* compared to larvae fed with ds*GFP* (Fig. 2c). Together with our observation that bacteria translocate to the hemocoel in *P. versicolora* larvae fed with insecticidal dsRNA, these findings indicate that the gut microbiota is an important contributor to dsRNA-mediated killing of pest insects.

A previous study showed that dsRNA-mediated gene knock-down in the migratory locust can induce overgrowth of opportunistic pathogens and intestinal atrophy (31), but the reasons remained unclear. To clarify how RNAi influences gut bacteria, we analyzed bacterial growth in *P. versicolora* after ingestion versus injection of insecticidal dsRNA. We found that the knock-down of an essential gene has limited relevance to the increase in gut bacterial abundance (Fig. 4), in that *Srp54k* knock-down triggered by injection of ds*Srp54k* had no statistically significant effect on total bacterial growth. Similarly, the increase in bacterial abundance in larvae injected with ds*Actin* was lower than that in larvae fed with ds*Actin*. Thus, the direct comparison between ingestion and injection indicates that ingestion was required for accelerated mortality (Fig. 4). This finding raises the interesting question why oral administration leads to proliferation of the gut microbiota. As bacteria do not possess an RNAi machinery and no dsRNA uptake systems have been found in bacteria (32), we hypothesized that the dsRNA itself cannot trigger overgrowth of gut bacteria. A dsRNA-degradating enzyme (dsRNase) in the gut was first identified and characterized in *Bombyx mori*, and related enzymes were subsequently found in almost all orders of insects, including the beetle family (Coleopteran). Differences in the expression levels and activities of intestinal dsRNases are likely, at least in part, responsible for the observed differences in susceptibility to RNAi between different groups of insects (2, 7, 33–36). In this work, we found that a gut dsRNase (dsRNase1) is chiefly response for dsRNA degradation in the digestive tract of *P. versicolora* (Fig. 5a-e). Moreover, we observed that ingestion of dsRNA upregulated *dsRNase1* expression (Fig. 5c). Suppressed expression of *dsRNase1* strongly reduced dsRNA degradation and decreased the abundance of gut bacteria (Fig. 5f-g), suggesting a contribution of *P. versicolora* dsRNase1 to growth control of gut bacteria. This finding also raised the interesting possibility that gut bacteria utilize the degradation products of dsRNAs as nutrient source (Fig. 5a). Evidence in support of this idea was supplied by bacterial growth assays that revealed that NTP mix, uridine and inosine, but not dsRNA, can be utilized as carbon and nitrogen sources to promote the growth of three dominant bacterial strains in the gut microbiome of *P. versicolora* (Fig. 5h). Thus, ingested dsRNA can suffer one of two fates in the gut of *P. versicolora* larvae: (i) it can be taken by midgut cells by a SID-type channel (putative dsRNA-selective transport channel) and/or the endocytosis pathway and subsequently trigger the RNAi response in the insect (37), or (ii) it undergoes degradation by dsRNase1 in the midgut (Fig. 5a) and its breakdown products are subsequently utilized by gut bacteria as carbon and nitrogen source for their growth (Figs. 5g and 7). However, it seems reasonable to assume that dsRNA is not be the only nutrient that controls gut bacterial growth and there are likely other factors that contribute to the observed gut bacterial dysbiosis.

**Fig. 7.**
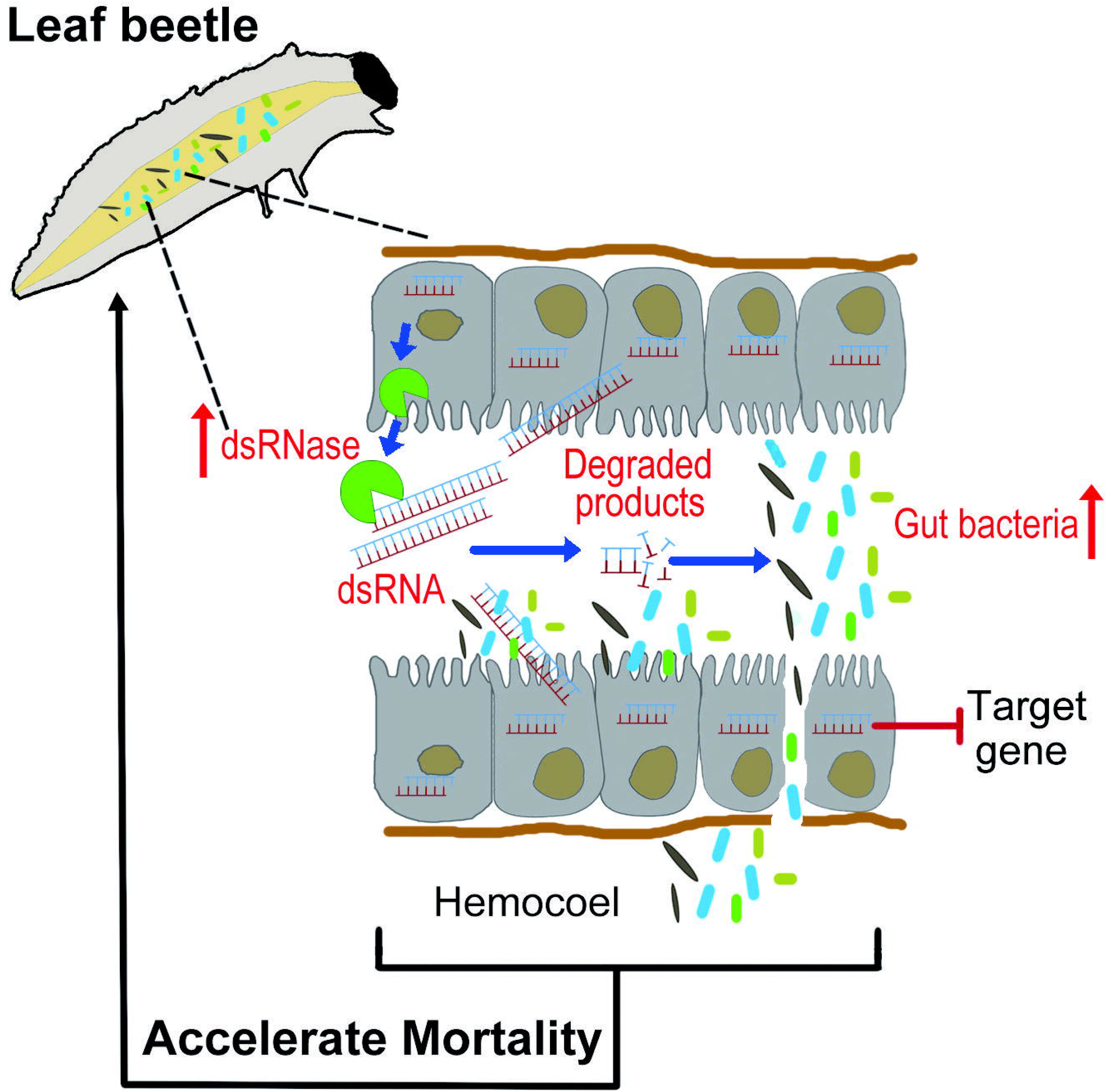
Working model of dsRNA-mediated insect killing. In the gut of the insect, dsRNA can be taken by midgut epithelial cells and loaded into the RNAi machinery, thus suppressing expression of the target gene. dsRNA can also induce expression of a gene encoding dsRNase, thereby causing degradation of dsRNA retained in the gut. Gut bacteria can utilize the degradation products of dsRNAs for their growth and translocate from the gut lumen into the hemocoel, thus resulting in accelerated mortality of their host insect.

It seems noteworthy that the capabililty of *P. putida* to utilize dsRNA degradation products as both carbon and nitrogen source for its growth is more pronounced than that of the other two abundant gut bacterial species tested (Fig. 5h; Table S3). This finding may, at least in part, explain why the proportion of *P. putida* was drastically increased in larvae fed with insecticidal dsRNAs (Fig. 2c). Moreover, re-introduction of gut bacteria into axenic larvae increased their mortality after feeding with lethal dsRNAs, and *P. putida* showed the strongest effect on acceleration of larval mortality (Fig. 6). *P. putida* was previously shown to be a nitrogen-fixing strain that can promote the growth of plants (38, 39), but whether the *P. putida* population in the gut microbiome of *P. versicolora* was originally acquired from its host plants and now functions as mildly pathogenic strain to *P. versicolora* remains to be investigated.

In conclusion, the present study illuminates the complex interplay between insecticidal dsRNA, the insect host, and its gut microbiota. Our work provides new insight into these interactions and suggests an important role of host sepsis in the multifaceted killing mechanism that underlies environmental RNAi.

## Acknowledgements

This work was supported by grants from the National Key Research and Development Program of China (2017YFD0600101), the National Natural Science Foundation of China (31572071, 31971663) and the Recruitment Program of Global Experts (China) to J.Z.

## Author contributions

LX and JZ conceived the project; LX, SX, LS, YZ, and JL performed the research; LX, SX, and JZ analysed the data. LX, RB and JZ wrote the article with contributions of all the authors.

